# Heme-NO dilates arteries via mobilization of NO moieties from a vascular intracellular NO store

**DOI:** 10.1101/2025.04.22.650128

**Authors:** Taiming Liu, Meijuan Zhang, Lingchao Zhu, Haiyan Ke, Amancio de Souza, Qian Li, Daniel Castella, Nicolai Lehnert, Lubo Zhang, Arlin B. Blood

**Affiliations:** Department of Pediatrics, Division of Neonatology, Loma Linda University School of Medicine, Loma Linda, CA 92354, USA; Department of Chemistry, University of California, Riverside, USA; Metabolomics Core Facility, University of California, Riverside, USA; Department of Medicine, Gregory Fleming James Cystic Fibrosis Research Center, University of Alabama at Birmingham, USA; Department of Chemistry, University of Michigan, Ann Arbor, MI 48109-1055, USA; Lawrence D. Longo, MD Center for Perinatal Biology, Loma Linda University School of Medicine, Loma Linda, CA 92354, USA

**Author notes:** Correspondence to: Taiming Liu. PhD, Department of Pediatrics, Division of Neonatology, Loma Linda University School of Medicine, Loma Linda, CA, Arlin Blood. PhD, Lawrence D. Longo, MD Center for Perinatal Biology, Loma Linda University School of Medicine, Loma Linda, CA.

**Keywords:** NO-ferroheme, nitrodilator, NANOS, intracellular NO store, thiyl radical

## Abstract

Nitrosyl heme (heme-NO) has recently emerged as a surrogate signaling entity for nitric oxide (NO). However, questions remain about how heme-NO signals across the cell membrane. Herein, we test the hypothesis that heme-NO signals as a nitrodilator that vasodilates by mobilizing NO moiety from a nitrodilator-activated intracellular NO store (NANOS) in the vasculature. We identify a novel mechanism for glutathione-catalyzed formation of a model compound, alb-heme-NO, and determine glutathione (GSH) as a ligand in its structure. Heme-NO complexes with plasma proteins in blood and, as such, it is impermeable to red blood cells or the vascular wall. Alb-heme-NO-mediated vasodilation, both *in vitro* and *in vivo*, is attenuated by prior depletion of the NANOS and potentiated by NANOS supplementation. Incubation with alb-heme-NO induces efflux of NO moieties from arteries. Additionally, the role of nitrosyl hemoglobin (HbNO) in mediating NO bioactivity export from erythrocytes is challenged. In conclusion, heme-NO functions as an extracellular nitrodilator via activation of the intracellular NANOS.

## Introduction

The canonical view of nitric oxide (NO) mediated vasodilatory signaling is that NO diffuses freely from endothelial NO synthase (NOS) to soluble guanylate cyclase (sGC) in vascular smooth muscle cells. However, since NO is a highly reactive radical with a short half-life, it remains questionable how NO survives various scavenging reactions to selectively activate sGC^1^. Several NO adducts, such as S-nitrosothiols (SNO) and dinitrosyl iron complexes (DNIC), have been proposed as intermediates that preserve the vasodilatory signaling potency of NO, although the mechanism by which these intermediates activate sGC is not yet characterized^2^. Nitrosyl heme (heme-NO) is another endogenous NO adduct that stabilizes NO and has been proposed as an intermediate for exporting NO bioactivity from red blood cells (RBCs)^3^. Unlike SNO and DNIC, heme-NO can irreversibly bind to and activate sGC as effectively as NO, potentially establishing it as a selective sGC agonist^4,5^. Recent studies propose that heme-NO acts as a surrogate signaling entity for NO^3,6,7^. However, a key uncertainty in this hypothesis is how heme-NO, as a bulky complex, signals as efficiently as NO across cell membranes^3,6^.

Previous work by us and others has revealed remarkable vasodilatory potency of SNOs and DNICs despite their membrane impermability and slow release of NO, raising the question: How do these NO adducts activate cytosolic sGC when their NO moieties are still outside the cell^8^? We defined these NO-containing and NO-mimetic compounds as nitrodilators, a concept that differs from nitrovasodilators that cause vasodilation via donation of free NO. We propose that nitrodilators induce vasodilation not by release of their own NO moieties, but rather by mobilizing NO moieties from a preformed NO store within vascular smooth muscle cells—referred to as the Nitrodilator-Activated intracellular NO Store (NANOS) model^2,8,9^. The NANOS is both depletable and repletable, corresponding to the attenuation and potentiation of nitrodilator-mediated vasodilation, respectively. This model proposes that the NANOS is regulated by the homeostasis of NO bioavailability. Our previous work suggested that nitrodilators, such as GSNO, DNIC, and the recently identified nitroglycerin (NTG), deplete NANOS in a use-dependent manner, while the NOS inhibitor L-NMMA depletes it by inhibiting endogenous NO production^2,8–10^. In contrast, despite being a NOS inhibitor, L-NAME unexpectedly contributes to the NANOS by slowly releasing NO from its nitro groups^11^. The enantiomer D-NAME, generally considered NOS-inactive and often used as a negative control for L-NAME, also contributes to the NANOS^11^. More importantly, nitrite—a circulating NO metabolite proposed to mediate the endocrine effects of NO and particularly abundant in the arterial wall—reverses the use-dependent tolerance of nitrodilators by replenishing the NANOS, potentially by increasing NO bioavailability^2,8–10^. The NANOS model represents a network of NO metabolites that function synergistically across cell membranes to facilitate NO signaling^2^.

A significant advancement in heme-NO research is the recent observation that glutathione (GSH) serves as a catalyst for the synthesis of heme-NO complexed with albumin, known as alb-heme-NO^3^. In the current study, we reexamine the role of GSH in alb-heme-NO synthesis and its impact on the structure of heme-NO. Using purified alb-heme-NO, we demonstrate that while heme-NO can be exchanged between albumin and other carriers, it remains membrane-impermeable and stable in blood. Next, we test the hypothesis that heme-NO signals across the cell membrane as a nitrodilator via mobilizing NO moiety from the NANOS. Additionally, we explore the role of heme-NO in exporting NO bioactivity from RBCs.

## Methodology

### Materials

5-tert-butoxycarbonyl-5-methyl-1-pyrroline-N-oxide (BMPO) was purchased from Enzo Life Sciences (Farmingdale, NY). ^15^N-nitrite and -nitrate were purchased from Cambridge Isotope Laboratories (Tewksbury, Ma). Nitroglycerin transdermal patch (Nitro-Dur^®^) was purchased from Mylan Pharmaceuticals (Canonsburg, PA). Glutathione (GSH; ^13^C_2_, ^15^N_1_, or unlabelled) and all other reagents were obtained from Sigma Aldrich (St Louis, MO). Sephadex G-25 columns were purchased from Cytiva Life Sciences (Marlborough, MA). NO gas was either sourced as a pure compressed gas (Matheson Tri-Gas, Inc, Irving, TX) or generated in a syringe by reacting sodium nitrite with sulfuric acid, then purified using deoxygenated NaOH. Whole blood (WB) was collected from adult ewes or 7 to 14-day-old lambs. Hemoglobin-NO (HbNO) in whole blood (WB) was prepared by reacting WB, deoxygenated by equilibration with N_2_ gas followed by addition of dithionite, with NO gas in the headspace of a syringe. The red blood cells of this WB were washed with deoxygenated saline by centrifugation, lysed by three freeze-thaw cycles, and further purified with a G-25 column to obtain free HbNO. For the lamb experiments, deoxygenated WB was reacted with NO gas to produce ∼50% HbNO, as monitored by spectrophotometry. Myoglobin-NO (MbNO) was prepared as reported^6^. Albumin (500 μM) was pretreated with N-ethylmaleimide (NEM; 5mM) and desalted using ultracentrifuge tubes with a 3 kDa cutoff (Amicon^®^, Millipore Sigma; Burlington, MA). Alb-heme-NO preparation, similar to previous report^3^, was achieved by reacting 500 μM albumin, 300 μM hemin (6 mM stock in 50 mM NaOH), 3 mM GSH, and 300 μM PROLI NONOate (10 mM stock in 10 mM NaOH) in HEPES buffer (final pH = 7.05±0.02) under anoxic conditions. The concentration of alb-heme-NO was calculated based on the concentration of hemin. Both MbNO and alb-heme-NO were further purified with G-25 columns at least once. S-nitroso-glutathione (GSNO) was synthesized as previously described^12^.

### Surgical preparations and infusion protocols

All procedures involving animals were performed according to the National Institutes of Health Guide for the Care and Use of Laboratory Animals, and were preapproved by the Loma Linda University (LLU) Institutional Animal Care and Use Committee (IACUC #21-198, #24-008, and lamb IACUC#8180027).

### Rat Protocol

Female non-pregnant Sprague-Dawley rats weighing 301±5 g were surgically instrumented as previously reported^9^. Rats were divided into several study groups. Each group received an i.p. injection daily for four days prior to and including the day of the experiment. The injectate was one of the following: 50 μmol/kg nitrite, 222 μmol/kg L-NAME, 222 μmol/kg D-NAME, 222 μmol/kg L-NMMA, or saline (Control). An additional group received a nitroglycerin (8 μg/h/kg) transdermal patch daily for four days. An extra group received only one i.v. injection of 222 μmol/kg L-NAME at 10 min prior to the administration of alb-heme-NO. Alb-heme-NO was administered either by i.v. infusion of 50 μM at stepwise increasing rates starting at 0.05 ml/min, then 0.1, 0.2, and 0.4 ml/min, with each rate maintained for 3 minutes, or as a bolus injection of 0.4 ml of a 300 μM solution into the jugular vein.

### Sheep Protocol

Lambs (Nebeker Ranch; Lancaster, CA) between 7 to 14 days of age were surgically instrumented, as previously described^13^. See the Supplemental materials for details.

### Wire myography

Sheep mesenteric arteries were dissected from isoflurane-anesthetized ewes, denuded of endothelium, and mounted in organ bath chambers as previously described^14^. For some experiments, the vessels were first exposed to three rounds of GSNO (5 μM) or UV light with each round lasting for 15 min. The vessels were then preconstricted for measurements of dilatory responses to alb-heme-NO. In some experiments, 10 μM 1H-[1,2,4]oxadiazolo[4,3,-a] quinoxalin-1-one (ODQ), 10 μM nitrite, 200 μM 2-(4-carboxyphenyl)-4,5-dihydro-4,4,5,5-tetramethyl-1H-imidazol-1-oxyl-3-oxide potassium salt (CPTIO), and/or 1000 U/ml superoxide dismutase 1 (SOD1) was added before tuning for basal tension.

See the Supplemental materials for details.

## Analytical methodologies

### Electron paramagnetic resonance (EPR) measurements

Unless specifically stated, EPR signals were recorded using a Bruker X-Band EMX Plus EPR spectrometer with a cavity of high sensitivity as previously described^15^. The EPR was set to a microwave power of 20 mW, microwave frequency of 9.34 GHz, attenuator of 10 dB, modulation amplitude of 1 G, modulation frequency of 100 kHz, time constant of 20.48 msec, conversion time of 81.92 msec, harmonic of 1, and number of scans of 2. For measurements of thiyl radical, using a previously described methodology^15^, BMPO (25 mM; spin trap), DMSO (500 mM; hydroxyl scavenger), and SOD1 (1000 U/ml;superoxide scavenger) were mixed in Hepes buffer with albumin (500 μM), hemin (300 μM dissolved in 50 mM NaOH), GSH (3 mM), and/or PROLI NONOate (300 μM) and measured at 6.0 min from the initiation of the reaction. Heme-NO, GSH-heme complexes, and arterial samples were measured at 110 K, while thiyl radical was measured at 293 K. Heme-NO concentrations were quantified using a standard curve of purified alb-heme-NO, unless otherwise specified. EPR signals of alb-heme-NO prepared with NaHS or nitrite (Figure S1f-g) were recorded using Bruker Magnettech ESR 5000 at 77 K with microwave power of 10 mW and microwave frequency of 9.50 GHz.

### Chemiluminesence measurements

The NOx (^14^N + ^15^N) levels were measured by a combination of five different assays with an ozone-based chemiluminescence NO analyzer (280i, Sievers, Boulder, CO) as previously described^16,17^. The selectivity of the five assays are given in Table S1 and supplemental materials.

### Gas Chromatography Mass Spectrometry (GC-MS) measurement

The ^15^N-NOx were measured under negative-ion chemical ionization mode by GC-MS (6890-5973; Agilent) as previously described^11^. See the Supplemental materials for details.

### Liquid Chromatography Mass Spectrometry (LC-MS) measurements

The measurements were performed on a G2-XS quadrupole time-of-flight mass spectrometer (Waters) coupled to an H-class UPLC system (Waters). See the Supplemental materials for details.

### Spectrophotometry measurements

The HbNO saturation was calculated via deconvolution of the UV‒ VIS spectrum by multiple linear regression analysis using basis spectra for HbO_2_, deoxyHb, metHb, HbNO, and HbCO as described before^18^.

## Results

### Role of GSH in its catalyzed formation of alb-heme-NO

We prepared alb-heme-NO as described by the previous study^3^, with two key modifications to eliminate high- and low-molecular-weight SNO byproducts^19^ before functional studies: albumin was pretreated with NEM to block thiol residues, and the alb-heme-NO products were purified using a G-25 column. While this study primarily focuses on the cross-membrane signaling mechanism of alb-heme-NO, several notable chemical observations that differ from the previous study^3^ are worth highlighting.

The addition of GSH to the reaction of albumin, hemin, and NO significantly enhanced the formation of alb-heme-NO (Figure 1a). However, the full EPR sweep of the reaction mixture also revealed rhombic signals, in addition to the canonical^20^ 5-coordinate (5-C) heme(Fe^2+^)-NO signal (g= 2.0725, 2.0298, and 2.0083) with a hyperfine coupling constant (A^N^) of 16.13 G (Figure 1b). These rhombic signals (g= 2.2758, 2.1854, and 1.9458) were partially reported in the previous study of alb-heme-NO product(s)^3^, and were similar to those simulated for GS-heme(Fe^3+^)-OH_2_ (g = 2.280, 2.158, 1.935; Figure S1a) and to those observed in 6-coordinate (6-C) thiolate-heme(Fe^3+^)-OH_2_ complexes, such as cytochrome P450^21^. These rhombic signals persisted even after G-25 column purification and were absent in HbNO and MbNO (Figure 1b; Refer to Figure S1 for comparisons of the low-spin components across various heme-NO complexes). These results indicated that our G-25 column-purified alb-heme-NO products contained both 5-C heme(Fe^2+^)-NO and an unidentified 6-C GS-heme(Fe^3+^) compound with rhombic signals in EPR. Notably, the unidentified 6-C GS-heme(Fe^3+^) compound is not 6-C GS-heme(Fe^3+^)-NO ({FeNO}^6^), as the latter is EPR silent, though it is also likely present in our alb-heme-NO products as will be discussed below. EPR quantification using an HbNO standard estimated that 27.1±2.2% (n=3 parallel measurements) of the heme in our alb-heme-NO products was heme(Fe^2+^)-NO. For clarity, the mixture of alb-heme-NO products is collectively referred to as alb-heme-NO throughout this manuscript.

**Figure 1.**
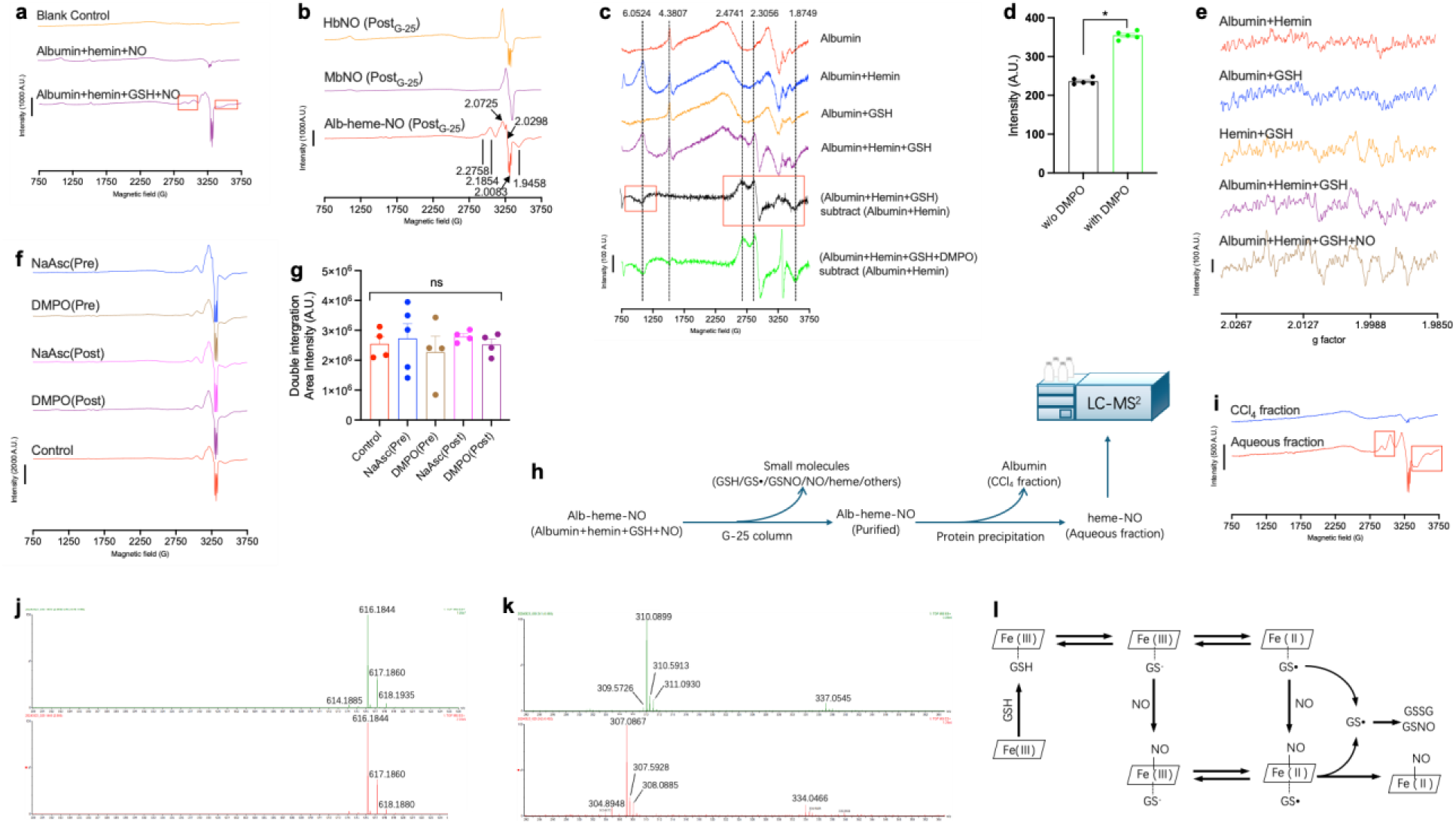
Role of GSH in alb-heme-NO synthesis and structure. n=4 to 5. **a** Representative EPR spectra of the reaction mixtures at 3 min of reaction. Note the highlighted (red box) additional rhombic signals around the heme-NO signal. **b** Comparison of the full scan EPR spectra of HbNO, MbNO, and alb-heme-NO post G-25 column purification. **c** Representative EPR spectra. **d** Comparison of the intensity of the EPR signals of GS-heme. Paired t-test. **c-d** GSH reduced high-spin ferric heme signal (left red box in **c**) and formed a low-spin GS-heme complex (right red box in **c**) in the absence of NO. The GS- heme complex formation was not blocked but rather enhanced by thiyl scavenger DMPO. **e** Thiyl radical was generated during the complexing of GSH and heme in the absence of NO, although NO facilitated the thiyl radical generation. **f** Representative EPR spectra. **g** Double integration quantification of the heme- NO signals in **f**. **f-g** The intensity of alb-heme-NO was not significantly altered by thiyl scavengers NaAsc and DMPO added before or after the synthesis. One-way ANOVA. **h** Protocol diagram of the MS experiments. Briefly, the synthesized alb-heme-NO was purified using a G-25 column to remove low molecular weight molecules, followed by albumin precipitation with CCl_4_ (1:1 volume ratio) before MS measurements of the aqueous fraction. **i** Representative EPR spectra confirming that the aqueous fraction retained the heme-NO after the CCl_4_ mediated albumin precipitation. Note the highlighted (red boxes) rhombic peaks persisted. **j-k** MS detection of heme (**j**; m/z=616) and GSH (**k**; m/z=307) in the aqueous fraction. **j-k** Bottom and top are results of alb-heme-NO synthesized with isotopically unlabeled and labeled (^13^C_2_, ^15^N_1_; m/z=310) GSH, respectively. See supplemental figure S3 for detailed MS spectra. **l** Proposed mechanism for the GSH-catalyzed formation of alb-heme-NO.

Thiols are known to bind ferric heme iron, forming thiolate-heme(Fe^3+^) complexes^15^. Consistent with this interaction, a full EPR sweep of the reaction mixtures (Figure 1c) showed that, in the absence of NO, GSH reduced the signal of high-spin ferric heme (g=6.0524), forming a low-spin GS-heme(Fe^3+^) complex (Figure 1c black line), which differ from the above rhombic signals observed in the presence of NO but are similar to those of the substrate-free thiolate-heme(Fe^3+^) cytochrome P450^22^. It has been proposed that the thiolate-heme(Fe^3+^) complex may equilibrate with thiyl radical-heme(Fe^2+^) via inner-sphere electron transfer^23^. Supporting this equilibration, the GS-heme(Fe^3+^) signal was enhanced by the thiyl radical scavenger DMPO (Figure 1d), possibly due to DMPO (EPR silent) converting GS•-heme(Fe^2+^) to DMPO-GS-heme(Fe^3+^).

A thiyl radical, similar to that previously reported^15^, was generated during the complexing of GSH and heme(Fe^3+^) in the absence of NO (Figure 1e), confirming that NO was not required for the aforementioned interaction of GSH and heme(Fe^3+^). Interestingly, NO further facilitated the thiyl radical generation, possibly by forming 6-C GS-heme(Fe^3+^)-NO and the GS•-heme(Fe^2+^)-NO intermediate, which may stabilize the thiyl radical^24^ that would otherwise cleave, releasing GS• and forming GSSG and GSNO. This facilitation was observed in the simple reaction of GSH and NO in the absence of albumin and heme (Figure S2), corroborating GSH as the source of the thiyl radical signal. Thiyl radical scavengers NaAsc and DMPO, added before the synthesis of alb-heme-NO, did not affect the yield of heme(Fe^2+^)-NO, and when added after synthesis, they did not affect its stability (Figure 1f-g). These results suggested that thiyl radical formation was either not essential or bypassed in the formation of heme(Fe^2+^)-NO.

Next, we used LC-MS (Figure 1h) to investigate the role of GSH in the structure of alb-heme-NO. For this investigation, the synthesized alb-heme-NO was purified using a G-25 column to remove low molecular weight molecules, followed by albumin removal through CCl_4_-mediated precipitation and phase separation. EPR measurements confirmed that the heme(Fe^2+^)-NO signal, along with the rhombic signals introduced by GSH, remained in the aqueous fraction after protein precipitation (Figure 1i).

Consistent with previous studies showing that albumin traps heme through its tyrosine residues^25^, these findings suggest that NEM-pretreated albumin, used as a heme solubilizer, is not a ligand in alb-heme-NO. More importantly, MS detected both heme and GSH in the aqueous fraction, a finding further confirmed by alb-heme-NO synthesized with isotopically labeled GSH (Figure 1j-k and Figure S3).

Although molecular ions of GS-heme or GS-heme-NO— fragile metal adducts in the ion source—were not detected, these results confirmed the presence of a GS- ligand in our alb-heme-NO products. The proposed mechanism for GSH-catalyzed alb-heme-NO formation is illustrated in Figure 1l. The chemical nature of the unidentified 6-C GS-heme(Fe^3+^) compound remains to be determined and is therefore not included in Figure 1l.

For the first time, alb-heme-NO was also prepared using NaHS in place of GSH and nitrite in place of NO, yielding final products with pH values of 7.63 ± 0.01 and 7.45 ± 0.01, respectively (Figure S1f-g). These findings extend the physiological relevance of heme-NO by demonstrating alternative pathways for its formation.

### Membrane permeability and stability of alb-heme-NO

We examined the permeability of heme-NO across the RBC membrane using plasma and whole blood (WB) as volume controls (Figure 2a-c). Compared to the volume controls, alb-heme-NO added to WB was nearly undetectable in the RBC fraction and instead concentrated in the plasma, suggesting that alb-heme-NO neither bound to nor crossed the RBC membrane in blood. In contrast, alb-heme-NO added to plasma-free blood (PFB; plasma replaced with saline) was detected in the RBC fraction but not in the saline fraction, indicating that heme-NO partitioned with RBCs in the absence of plasma. When plasma was added back to these RBCs, it extracted most (76.08±2.57 %) of the heme-NO that had partitioned with RBCs (Figure 2d-f), suggesting that in the absence of plasma, heme-NO likely binds to the exterior of the RBC membrane rather than entering the cells. These results demonstrated that heme-NO is impermeable to the RBC membrane. They also suggested that heme-NO, which can bind to the RBC membrane, is preferably retained by certain plasma protein(s) in the blood. Consistent with these *in vitro* findings, measurements of plasma and RBC samples collected from rats before and after i.v. alb-heme-NO infusion revealed that the infused NOx was primarily retained in the plasma rather than the RBCs (Figure 2g).

**Figure 2.**
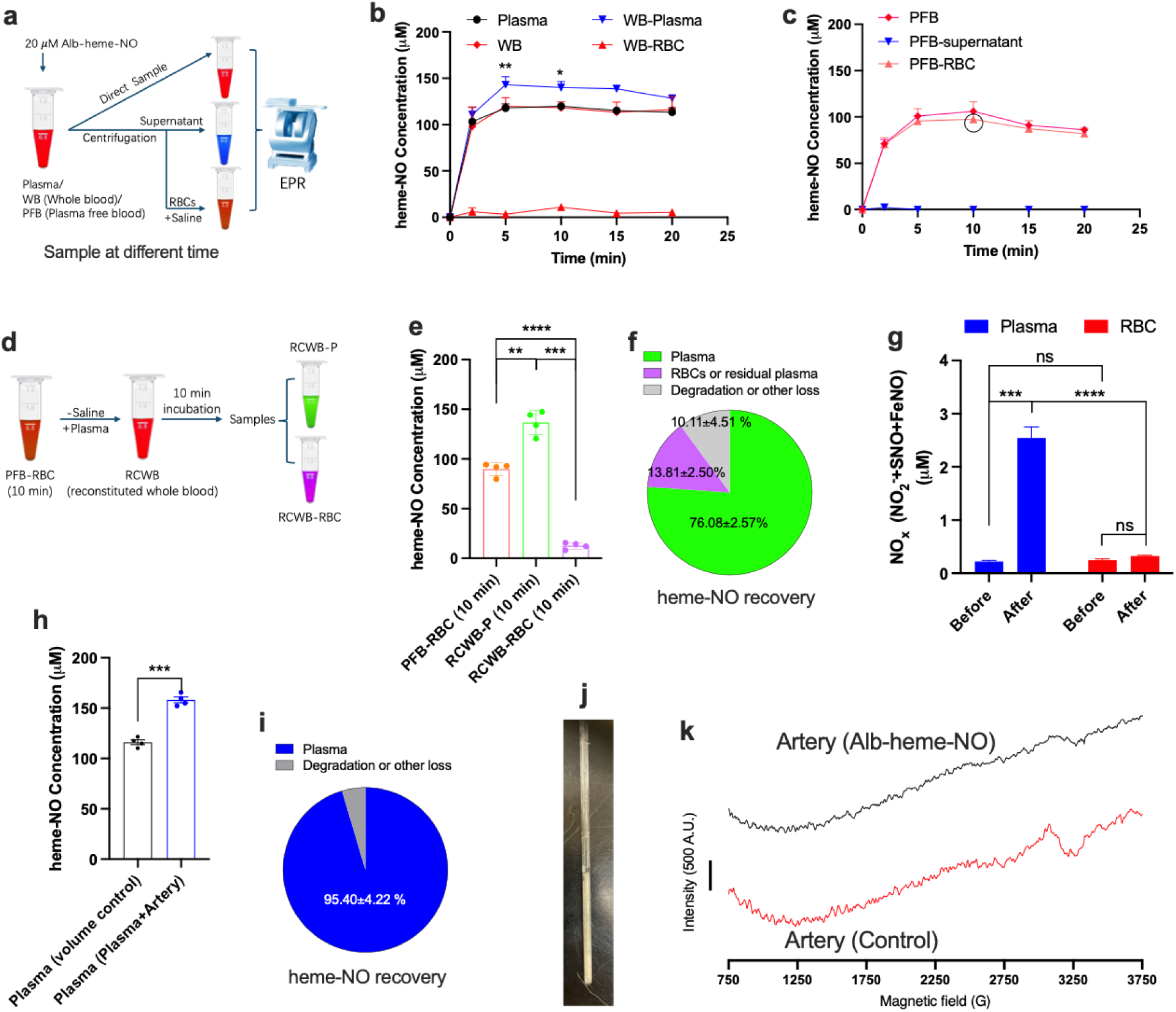
Membrane permeability and stability of alb-heme-NO. n=4 to 5. **a** Protocol diagram for **b** and **c**. Briefly, 20 μM G-25 column purified alb-heme-NO was added into plasma, whole blood (WB), or plasma free blood (PFB), then sampled at different time points either directly or after fractionation by centrifugation. Heme-NO was quantified by EPR using a standard curve of alb-heme-NO prepared in water. **b-c** Pharmacokinetics of alb-heme-NO in plasma and different fractions of WB (**b**) and PFB (**c**). Note that heme-NO was retained in the plasma of WB (Hematocrit= 70-75 %), but partitioned with RBCs in PFB. p-value refers to the result of paired t-test versus Plasma. **d** Protocol diagram for **e-f**. Briefly, saline in the PFB-RBC samples (prepared as those highlighted in the black circle of (**c**) was replaced with plasma to make reconstituted whole blood (RCWB; Hematocrit= 50 %). The RCWB was further incubated for 10 min before fractionation for EPR measurements. **e** EPR quantification using a standard curve of alb-heme-NO prepared in water. **f** Recovery of heme-NO calculated from **e**. Most heme-NO partitioned with PFB-RBC was extracted by plasma. p-value refers to the result of One-way ANOVA with Tukey’s post hoc test. **g** I_3_^-^-based chemiluminescence measurements of plasma and RBC collected from rats before and at the end of i.v. infusion of alb-heme-NO (Figure 4a). Paired t-test. **h-k** Membrane permeability of alb-heme-NO in isolated arteries in the presence of plasma. 20 μM alb-heme-NO was added into either a mixture of plasma and femoral artery (volume/weight ratio of 7:3) or plasma alone (as control), and then incubated for 20 min before sampling for EPR measurements. **h** EPR quantification of heme-NO in plasma fraction. Paired t-test. **i** Recovery of heme-NO calculated from **h**. Most heme-NO was retained by plasma. **j** Image showing 40 cm of sheep femoral arteries packed into 5 cm of an EPR tube. **k** Representative EPR spectra of arteries after 20 min incubation with 20 μM alb-heme-NO in plasma (top) and with plasma alone (bottom). Heme-NO was not detected in arteries in either case. All incubations were performed under 37 °C in dark.

Notably, the heme-NO concentrations in plasma, WB, and PFB increased over the first 5 minutes following alb-heme-NO addition, rather than peaking at the initial measurement (Figure 2b-c). These heme-NO concentrations in plasma, WB, and PFB peaked at around 100 μM when quantified using a standard curve of purified alb-heme-NO in buffer, far exceeding the initially added 20 μM. Additionally, EPR measurements showed that the rhombic signals assigned to the unidentified 6-C GS-heme(Fe^3+^) compound were no longer present in these samples (Figure S4-6). Finally, the plateaued kinetic curves of heme-NO in plasma, WB, and PFB suggested that heme-NO was stable in both plasma and blood.

We also examined the permeability of heme-NO across arterial membranes. To mimic *in vivo* conditions, alb-heme-NO was added to a mixture of plasma and isolated arteries, with plasma alone serving as the volume control. Heme-NO was found to be retained in the plasma of the mixture and was undetectable in the arteries (Figure 2h-k). These results are consistent with the above observations in blood, further confirming that heme-NO is cell membrane impermeable.

### *In vitro* functional evidence supporting the involvement of a nitrodilator activated NO store (NANOS) in alb-heme-NO mediated vasodilation

Consistent with a recent report^6^, the sGC oxidizer ODQ completely blocked alb-heme-NO-mediated vasodilation (Figure 3a). Despite its membrane impermeability, alb-heme-NO exhibited a relaxation dose-response curve with an logEC_50_ similar to that of free NO (p=0.2430 for -5.96 ± 0.21 vs -6.20 ± 0.24; Figure 3b), suggesting it may act as a vasodilator by mobilizing NO moieties from an intracellular NO store, i.e. the NANOS, preformed within vascular muscle cells. Supporting this notion, alb-heme-NO-mediated vasodilation was attenuated by pretreatment of the vessels with GSNO (Figure 3c-d), which has been previously shown to deplete the NANOS. Additionally, this attenuation was reversed by nitrite that has been shown to replenish the NANOS. Moreover, our recent study suggested that the NANOS overlaps with the intracellular NO store that can be depleted by exposure to UV light^8^. Consistent with this, UV pretreatment of vessels to deplete the NO store significantly attenuated alb-heme-NO-mediated vasodilation (Figure 3e). Similar to previously identified nitrodilators^2^, vasodilation mediated by alb-heme-NO was not affected by CPTIO or SOD1 alone, but was significantly attenuated when both were used in combination (Figure 3f). These findings provide evidence that alb-heme-NO is a nitrodilator that causes vasodilation through the mobilization of the NANOS (Figure 3g).

**Figure 3.**
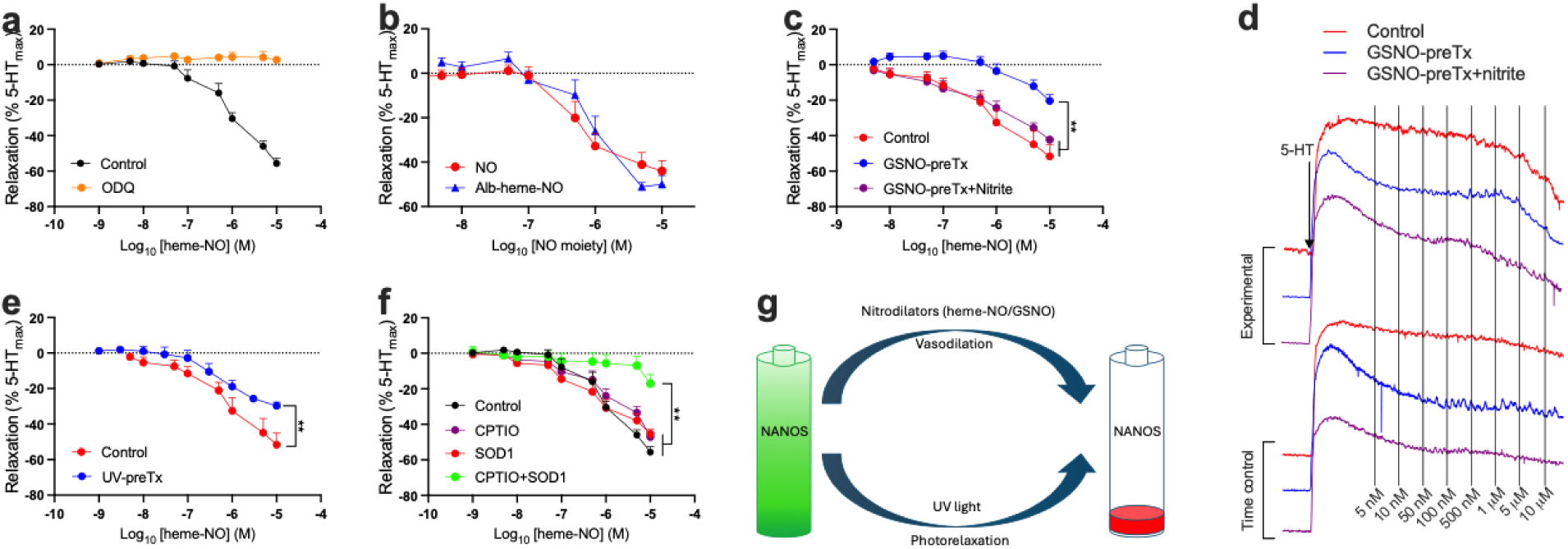
*In vitro* functional evidence supporting the involvement of NANOS in alb-heme-NO mediated vasodilation. *n*=5. **a** Effects of sGC oxidizer ODQ on alb-heme-NO-mediated vasodilation. **b** Comparable vasodilation mediated by alb-heme-NO and NO in isolated arteries. PROLI NONOate was used as the NO donor. **c-d** GSNO pretreatment (preTx) attenuated subsequent alb-heme-NO mediated vasodilation, an effect that was restored by nitrite. GSNO pretreatment was previously shown to deplete the NANOS, while nitrite was shown to replenish the NANOS^8,9^. **d** Representative traces of wire myography experiments in **c**. **e** UV pretreatment, previously shown to deplete the NANOS^8^, attenuated subsequent alb-heme-NO-mediated vasodilation. **f** Like other nitrodilators (GSNO, nitroglycerin, DNICs)^2,8,14^, alb-heme-NO-mediated vasodilation was not affected by CPTIO (NO scavenger) or SOD1 (converts HNO into NO) alone, but significantly attenuated by their combination. **g** Proposed diagram for the role of the NANOS in alb-heme-NO-mediated vasodilation. We propose alb-heme-NO is a member of nitrodilators. All wire myography experiments were conducted using endothelium-denuded sheep mesenteric arteries. p-value refers the result of paired t-test versus Control.

### *In vivo* functional evidence of alb-heme-NO-mediated vasodilation through NANOS activation

We next examined the role of the NANOS in alb-heme-NO-mediated vasodilation *in vivo*. As shown in Figure 4a-b, rats were pretreated with either a NANOS depleter (L-NMMA or NTG) or repleter (L-NAME, D-NAME, or nitrite)^2,8^ for four days before receiving a stepwise infusion or bolus injection of alb-heme-NO. During stepwise infusion, alb-heme-NO-mediated vasodilation in mesenteric arteries was attenuated by the NANOS depleters and enhanced by the NANOS repleters (Figure 4c). These NANOS modulators did not affect the pharmacokinetics of alb-heme-NO in plasma (Figure 4d). Consistent with the above *in vitro* observations (Figure 3), these results suggest that alb-heme-NO-mediated vasodilation *in vivo* also involves the NANOS. The effects of stepwise alb-heme-NO infusion on mean arterial blood pressure (MAP) and heart rate are shown in Figure S7.

**Figure 4.**
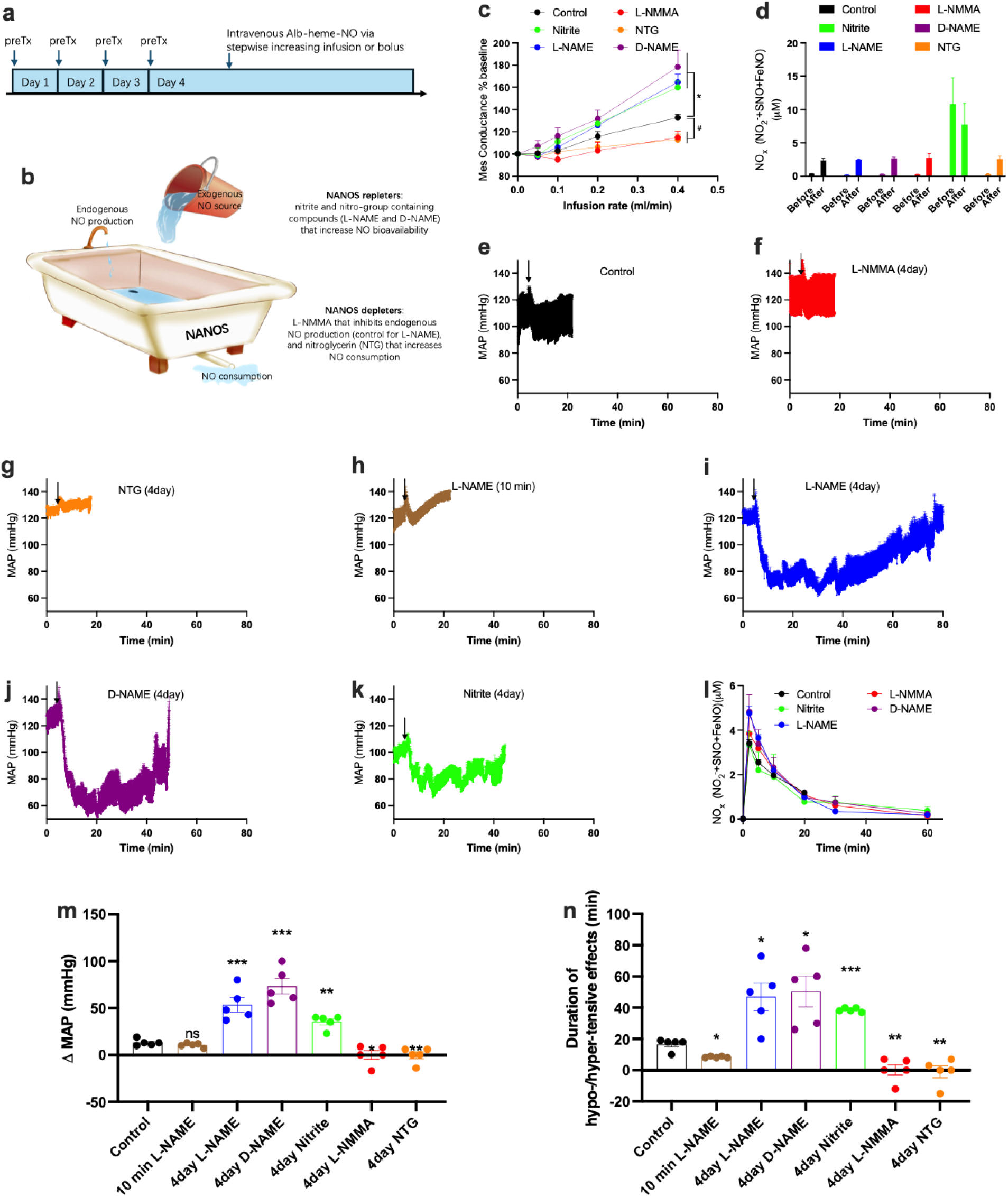
*In vivo* functional evidence supporting the involvement of NANOS in alb-heme-NO mediated vasodilation. *n*=5. **a** Protocol. Rats were pretreated (preTx) for 4 days with i.p. injection of a NANOS depleter or repleter. On day 4, alb-heme-NO was administered either by stepwise infusion or bolus injection into the jugular vein. **b** Diagram depicting the homeostasis of the NANOS: the NANOS is constitutively supplemented by endogenously generated NO and reduced by endogenous nitrodilators. Nitrite, L-NAME, and D-NAME were previously shown to increase the NANOS, while L-NMMA and NTG were previously shown to deplete the NANOS^2,8,11^. **c** The vasodilation in rat mesenteric arteries mediated by stepwise infusion of alb-heme-NO was potentiated by NANOS repleters nitrite, L-NAME, and D-NAME but attenuated by NANOS depleters L-NMMA and NTG. Two-way ANOVA. **d** I_3_^-^-based chemiluminescence measurements of total NOx in plasma before and at the end of alb-heme-NO stepwise infusion. **e-k** Mean arterial pressure (MAP) traces of rats upon bolus injection of alb-heme-NO (depicted by black arrow). **e** Control. **f** L-NMMA (4 days). **g** NTG (4 days). **h** L-NAME (10 min). In contrast with other pretreatments that were given for 4 days, in this group i.v. L-NAME was injected at 10 min prior to the alb-heme-NO bolus injection. **i** L-NAME (4 days). **j** D-NAME (4 days). **k** Nitrite (4 days). **l** I_3_^-^-based chemiluminescence measurements of total NOx in plasma after alb-heme-NO bolus injection. Baseline was corrected by subtraction of the NOx level measured before alb-heme-NO bolus injection. **m-n** Effects of NANOS modulators on the amplitude (**m**) and duration (**n**) of changes in MAP following intrajugular boluses of alb-heme-NO. Negative values represent opposite changes. p-value refers the result of unpaired t-test versus Control.

When injected as a bolus in control rats, alb-heme-NO caused a maximum decrease in MAP of 13.2±1.6 mmHg, which lasted for 16.3±2.1 minutes. NANOS depleters L-NMMA and NTG completely blocked the hypotensive effects of alb-heme-NO, while NANOS repleters nitrite, L-NAME, and D-NAME increased both the amplitude and duration of its hypotensive effects (Figure 4e-n). None of the NANOS modulators affected the pharmacokinetics of alb-heme-NO in plasma (Figure 4i). In an additional group of rats, administering L-NAME 10 minutes before alb-heme-NO did not enhance the hypotensive effects but instead shortened their duration (Figure 4h, m-n), consistent with previous work showing that the contribution of L-NAME to the NANOS is slower than its NOS-inhibiting effects^11^.

### Alb-heme-NO facilitates efflux of NO moieties from the arterial smooth muscle

Utilizing a recently established experimental model^8^, we next explored the chemical evidence supporting the role of the NANOS in alb-heme-NO-mediated vasodilation. Briefly, two sheep carotid arteries were first incubated with ^15^N-nitrite to load the NANOS with ^15^N (Figure 5a-b). Next, the arteries were sealed and incubated in parallel with either alb-heme-^14^NO in plasma to activate NANOS and release ^15^N-NO moieties, or plasma (vehicle) alone injected into the sealed lumen. Each artery was immersed in a buffer containing metHb on the abluminal side to capture the ^15^N-NO moieties released by NANOS. Both chemiluminescence and GC-MS measurements showed that alb-heme-NO decreased concentrations of an ^15^N-containing compound in the arterial wall which was detected as nitrate, and increased ^15^N-NOx levels in the abluminal buffer (Figure 5c-j), indicating that alb-heme-NO activated the efflux of ^15^N-NO moieties from the arterial wall. The proposed NANOS model is illustrated in Figure 5k.

**Figure 5.**
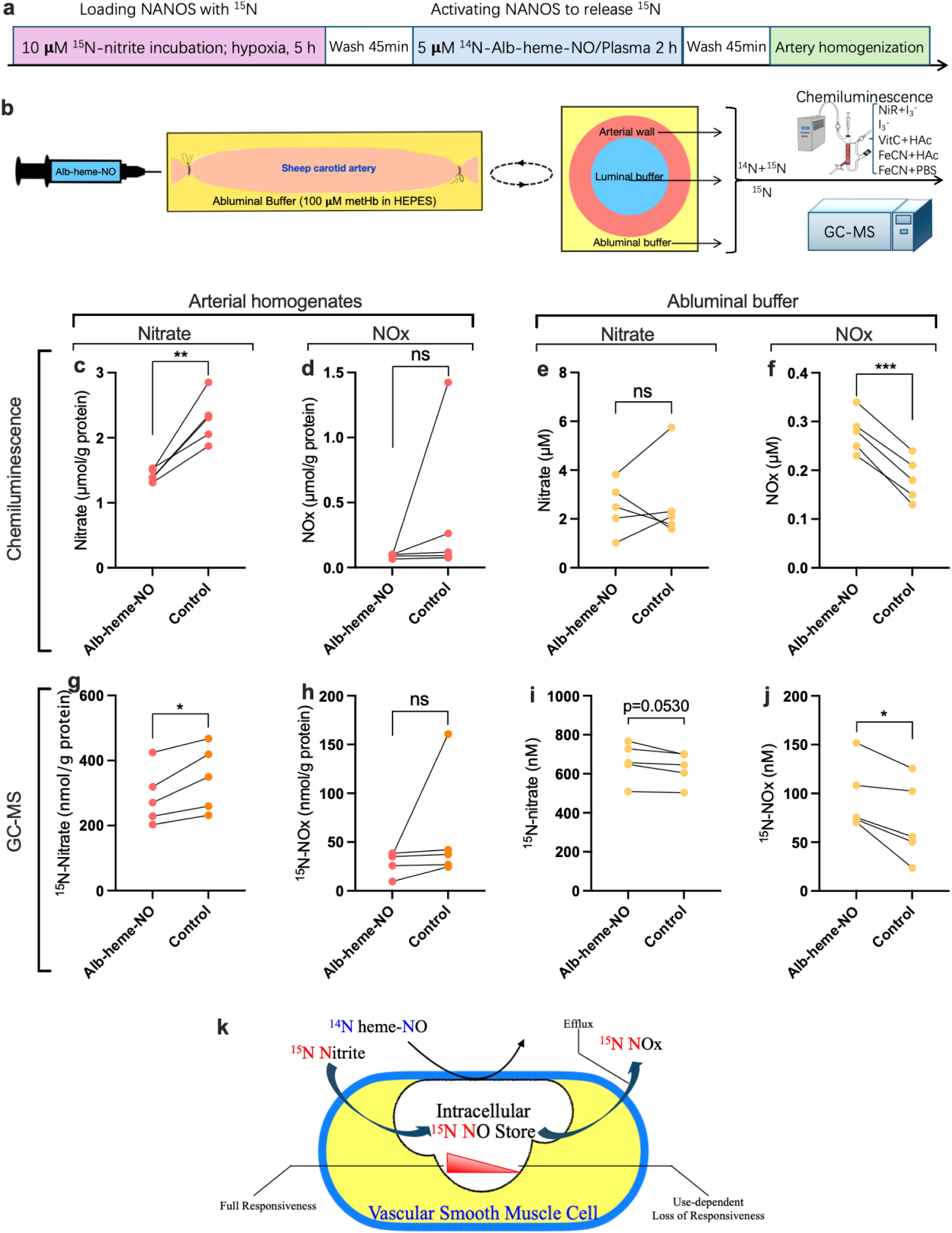
Alb-heme-NO facilitates efflux of NO moieties from the arterial smooth muscle. **a-b** Protocol. Sheep carotid arteries (length 8 cm; endothelium-denuded) were incubated with 10 μM ^15^N- nitrite in DMEM under 10.5% O_2_/5% CO_2_ for 5 h to isotopically load the NANOS with ^15^N-NO. After extensive washes, as depicted in **b**, the arteries were sealed and incubated for 2 h with 3 ml of 100 μM MetHb in HEPES buffer on the abluminal side and 5 μM alb-heme-NO or control (0.7 ml plasma) on the luminal side. Abluminal and luminal buffers as well as arterial homogenates were then assayed for NO species with five chemiluminescence assays and GC-MS. **c-f** Chemiluminescence measurements. **g-j** GC-MS measurements. **c, d, g, h** Measurements of arterial homogenates. **e, f, i, j** Measurements of abluminal buffer. Chemiluminescence measurements of nitrate were based on the use of nitrate reductase (NiR) + I_3_^-^, while that of NOx were based on the use of I_3_^-^. NOx includes nitrite, SNOs, and FeNO, but not nitrate. Alb-heme-NO stimulated an efflux of NO moieties from the arterial wall, providing chemical evidence for the NANOS hypothesis. Results of the other three chemiluminescence assays and all measurements of the luminal and control buffers are given in supplemental figures S8-12. **k** Diagram of the proposed NANOS model. Nitrodilator heme-NO causes vasodilation via mobilization of the NO moiety from a depletable and repletable NANOS in the artery. Note: N in red and blue denotes the different sources of the NO moieties. p-value refers the result of paired t test.

### Role of heme-NO in the export of NO bioactivity from RBCs

We also investigated the vasoactivity of various other heme-NO complexes. In wire myography (Figure 6a), the vasodilatory activity of alb-heme-NO prepared with dithionite, as previously reported^3^, was comparable to that of alb-heme-NO prepared with GSH. MbNO was significantly less potent than alb-heme-NO, while HbNO exhibited no vasoactivity, even under hypoxic or anoxic conditions (not shown).

**Figure 6.**
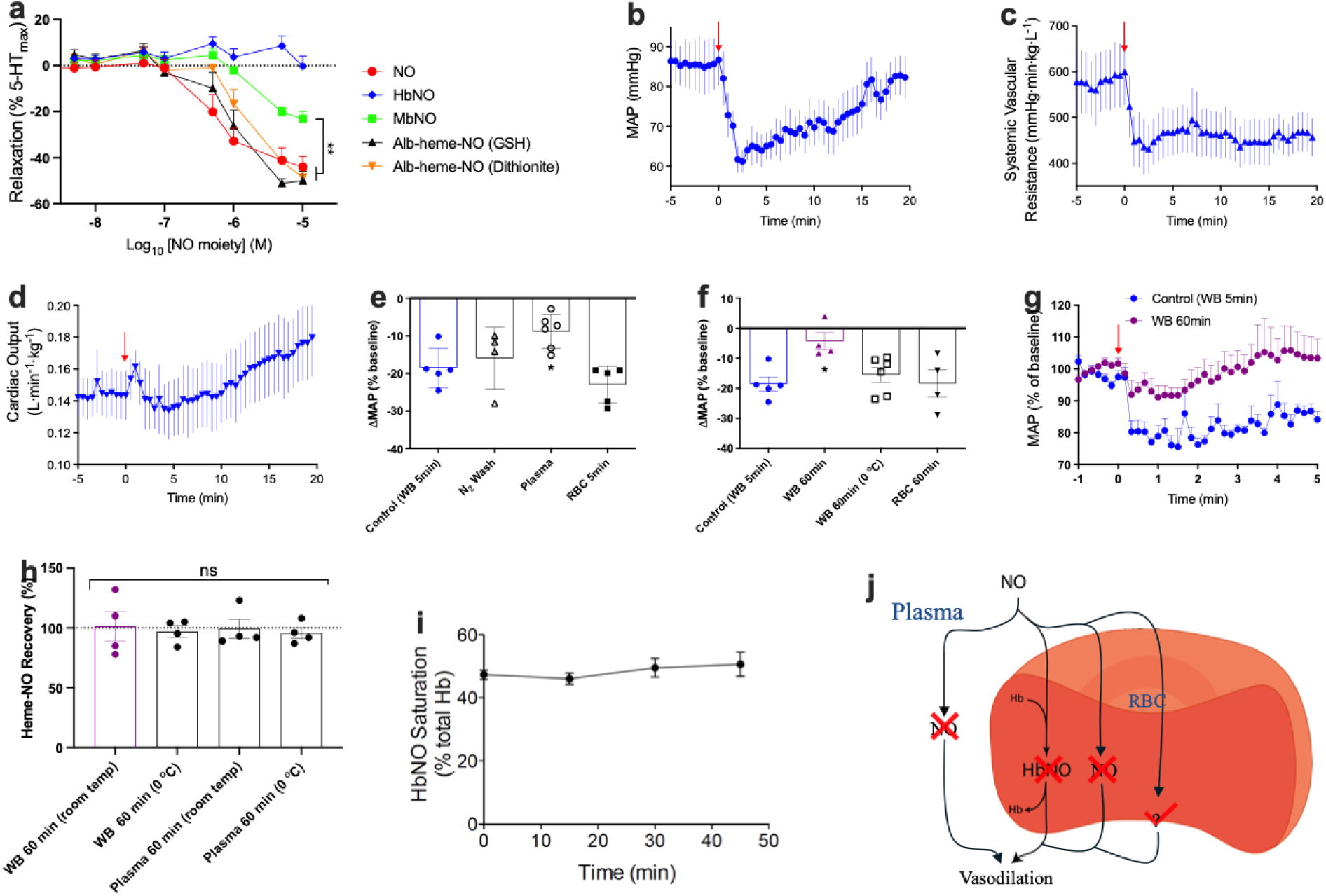
Role of heme-NO in the export of NO bioactivity from RBCs. **a** Comparison between the vasodilatory activity of NO and various heme-NO complexes including HbNO, MbNO, and alb-heme-NO prepared with GSH or dithionite. All heme-NO were purified with G-25 columns. n=5. **b-d** Bolus injection of 80 μM HbNO in adult whole blood resulted in decreases in MAP (**b**) and systemic vascular resistance (**c**), and a mild increase in cardiac output in lambs (**d**). **e** Comparison between the hypotensive effects of 20 μM fresh HbNO in lamb whole blood (WB 5 min; Control), Control washed with N_2_ gas (N_2_ wash), plasma saturated with NO (Plasma), and 20 μM fresh HbNO in RBCs (RBC 5 min) in lambs. * p<0.05 for unpaired t-test versus Control. **f** Comparison of the hypotensive effects of Control (WB 5 min), Control placed at room temperature for 60 min (WB 60 min), Control placed at 0°C for 60 min (WB 60 min (0 °C)), and RBC 5 min placed at room temperature for 60 min (RBC 60 min) in lambs. * p<0.05 for unpaired t-test versus Control. **g** Averaged MAP traces of Control (WB 5 min) and WB 60 min. Bolus injection of Control (WB 5 min) but not Control placed at room temperature for 60 min (WB 60 min) resulted in hypotensive effects. **h** EPR-measured stability of HbNO in whole blood (left two bars) and cell-free HbNO in plasma (right two bars) over a 60-minute period at room temperature or 0°C. One-way ANOVA. **i** Spectrometry-measured stability of HbNO in whole blood (Control sample/injectate under room temperature). **j** Proposed diagram for how whole blood pretreated with NO mediates vasodilation.

Given the potential relevance of HbNO in neonates undergoing inhaled NO therapy^26,27^, we further investigated the vasoactivity of HbNO in lambs. In contrast to the lack of effect observed with purified HbNO in wire myography (Figure 6a), HbNO in adult sheep whole blood (freshly prepared and used within 5 minutes) significantly decreased MAP and systemic vascular resistance while mildly increasing cardiac output in lambs. To explore the cause of this discrepancy, we compared the vasoactivity of HbNO in WB after various manipulations prior to injecting it into the lambs. First, equilibration of the sample with nitrogen gas did not significantly diminish the hypotensive effects of HbNO in WB, indicating that dissolved NO in the blood was not responsible for the observed hypotensive effects. Furthermore, plasma that was equilibrated with NO gas in a method similar to that used to generate HbNO in the whole blood samples produced significantly less hypotensive effects than HbNO in WB, ruling out the possibility that the hypotensive mediator resides in the plasma. Consistent with this, HbNO formed in plasma-free RBCs exhibited hypotensive effects similar to those of HbNO in WB, confirming that the hypotensive mediator resides within the RBC.

We next investigated the stability of the hypotensive mediator. HbNO in WB was significantly less hypotensive after 60 minutes of storage at room temperature, but not at 0°C. These results suggest that the hypotensive mediator was degraded during the 60 minutes at room temperature via a temperature-dependent pathway. However, HbNO in RBC remained hypotensive after 60 minutes of storage at room temperature, suggesting that the degradation of the hypotensive mediator involves components in the plasma. Importantly, EPR measurements demonstrated that heme-NO was stable in WB and plasma.

Additionally, spectrophotometry measurements showed that HbNO saturation in WB did not significantly decrease over 60 minutes. These results suggested that the hypotensive mediator is unlikely to be heme-NO or HbNO. The proposed diagram illustrating the role of heme-NO/HbNO in vasodilation mediated by WB pretreated with NO is shown in Figure 6j. It remains uncertain what serves as the hypotensive mediator.

## Discussion

This study identifies a novel mechanism for GSH-catalyzed formation of alb-heme-NO, and reveals GSH’s role as a ligand in the structure of alb-heme-NO. In contrast to the idea that heme-NO traverses the cell membrane for signaling, our experiments demonstrate that heme-NO is retained by plasma protein(s) and is impermeable to RBCs and the vascular wall. In addition, the role of HbNO as a source vasodilatory heme-NO exported from RBCs is challenged. More importantly, the current work provides both functional and chemical evidence that heme-NO causes vasodilation by mobilizing NO moieties from a vascular intracellular NO store.

### Catalytic role of GSH in alb-heme-NO synthesis

This study confirms the previously reported catalytic role of GSH in the formation of alb-heme-NO^3^. However, in contrast to the previous report identifying a reductive nitrosylation pathway, the current study reveals that GSH binds to heme(Fe^3+^) before complexing with NO. The discrepancy may stem from the fact that the current study was conducted in an aqueous buffer while the previous mechanistic investigation used methanol, which could facilitate reductive nitrosylation^28^. It is important to note that, since the reductive nitrosylation mechanism is not investigated in the current study, its existence cannot be ruled out. For the first time, this work directly detects thiyl radical formation in the reaction mixture of GSH and heme(Fe^3+^), with NO facilitating this process. Consistent with this observation, thiyl radical formation from GSH is facilitated by NO in the absence of heme(Fe^3+^). It is noteworthy that these observations of facilitation are unexpected, as thiyl radicals are thought to be diminished by their extremely rapid reaction with NO radicals to generate SNO^29^. The chemical implications of these findings warrant further investigation.

### Impacts of GSH on structure of heme-NO

In addition to 5-C heme(Fe^2+^)-NO, our alb-heme-NO products contained an unidentified 6-C GS-heme(Fe^3+^) compound, and likely also 6-C GS-heme(Fe^3+^)-NO. The identification GS-as an axial ligand bound to the heme-NO species in this work aligns with the proposition that GS-is such a highly favored ligand for heme that it has been suggested nearly all cytoplasmic heme exists in association with GS-^30^. Notably, it has been reported that 5-C heme(Fe^2+^)-NO exhibits g factors similar to those of 6-C thiolate-heme(Fe^2+^)-NO, attributed to the weakness of the S-Fe bond in thiolate-heme(Fe^2+^)^31^. Therefore, it remains possible that the 5-C heme(Fe^2+^)-NO measured by EPR in both the current and previous study^3^ was actually 6-C GSH-heme(Fe^2+^)-NO, with GSH loosely binding to the sixth coordination site of heme.

### Membrane permeability and kinetics of alb-heme-NO in blood

Heme-NO is an agonist of sGC and can exchange between different carrier proteins^4^. Since heme-NO is a potent vasodilator via activation of intracellular sGC without releasing significant amounts of free NO, it seems intuitive that it crosses the cell membrane to mediate signaling^3,7^. Contrary to this intuition, our EPR experiments on heme-NO partitioning in blood demonstrate that heme-NO is retained in plasma as opposed to entering RBCs and the vascular wall, suggesting it does not activate intracellular sGC directly. Intriguingly, upon the addition of alb-heme-NO to plasma, blood, and PFB, the rhombic signals of the unidentified 6-C GS-heme(Fe^3+^) compound disappeared, while those of 5-C heme(Fe^2+^)-NO increased.

These observations suggest that the unidentified 6-C GS-heme(Fe^3+^) compound and/or the putative EPR-silent 6-C GS-heme(Fe^3+^)-NO was efficiently converted into 5-C heme(Fe^2+^)-NO. The mechanism underlying the heme reduction of this conversion is beyond the focus of the current study and warrants further investigation. In addition to this conversion, several aspects of carrier protein exchange following the addition of alb-heme-NO to plasma, blood, and PFB merit noting. When heme-NO partitions with PFB-RBC in the absence of plasma, it is likely that the albumin in alb-heme-NO is exchanged by the lipid/protein components of the plasma membrane of RBC. However, when heme-NO partitions in the presence of plasma, it is retained by plasma protein(s). It remains to be determined how the exchange of the carrier occurs in plasma. The exchange of albumin by hemopexin, which has the highest known affinity for binding heme among all proteins, is less likely; otherwise, this would have produced an EPR signal characteristic of 6-C heme(Fe^2+^)-NO with proximal histidine as the axial N-donor ligand^25^.

### Wire myography evidence for the involvement of NANOS in heme-NO-mediated vasodilation

In this study, multiple lines of evidence from wire myography support the involvement of the NANOS in heme-NO-mediated vasodilation. First, alb-heme-NO mediated sGC-dependent vasodilation at concentrations comparable to that of NO, and the NO scavenger CPTIO did not block this effect, indicating that alb-heme-NO does not vasodilate via release of NO. As a bulky protein complex, alb-heme-NO cannot cross the plasma membrane, let alone with the efficiency of NO. Even though the heme-NO core of alb-heme-NO is exchangeable and may bind to the exterior of the plasma membrane in the absence of plasma, studies have demonstrated that the transfer of heme-NO is unidirectional, from the plasma membrane to albumin^3^. Therefore, it is unlikely that alb-heme-NO mediates vasodilation by directly activating intracellular sGC via its NO or heme-NO components, suggesting the presence of alternative sources of NO bioactivity for its activation of sGC. Second, it has previously been shown that pretreatment of vessels with UV light and GSNO induces tolerance in nitrodilator-mediated vasodilation by depleting the arterial NANOS^8,9^. Consistent with this, alb-heme-NO-mediated vasodilation was attenuated in vessels pretreated with UV light or GSNO, treatments that do not affect NO-mediated vasodilation^8,9^. More importantly, the attenuation of alb-heme-NO-mediated vasodilation by GSNO pretreatment was reversed by nitrite, which has been proposed to restore the NANOS level^8^. Third, our previous studies have suggested that the NO moieties released from the NANOS exhibit HNO-like characteristics, leading to nitrodilator-mediated vasodilation being unaffected by CPTIO (an NO scavenger) or SOD1 (which converts HNO into NO) alone, but significantly attenuated when both are used in combination^2,8,14^. Consistent with these observations, CPTIO and SOD1 had similar effects on alb-heme-NO-mediated vasodilation in the current experiments.

### *In vivo* evidence for the involvement of NANOS in heme-NO-mediated vasodilation

Consistent with the above *in vitro* evidence, our *in vivo* experiments also support alb-heme-NO being a nitrodilator. Vasodilation mediated by continuous infusion of alb-heme-NO in rats was attenuated by four-day pretreatments with an NTG patch and L-NMMA, which deplete the NANOS, and potentiated by four-day pretreatments with nitrite, L-NAME, and D-NAME, which contribute to the NANOS. Similarly, both the amplitude and duration of vasodilation mediated by bolus injection of alb-heme-NO were altered in accordance with these NANOS manipulations. Measurements of NOx in plasma following both continuous and bolus infusion of alb-heme-NO indicated that its pharmacokinetics were not significantly affected by the pretreatments. These results are consistent with the involvement of the NANOS in alb-heme-NO mediated vasodilation *in vivo*. Notably, L-NAME is expected to exert dual effects on the NANOS. On one hand, as evidenced by its rapid induction of hypertension within several minutes, L-

NAME is capable of swiftly inhibiting *in vivo* NO production, potentially reducing endogneous replenishment of the NANOS. On the other hand, L-NAME contributes to the NANOS by gradually, over a matter of hours, releasing NO from its nitro groups^11^. Therefore, short-term pretreatment of rats with L-NAME is expected to produce effects opposite to those of long-term pretreatment on alb-heme-NO mediated vasodilation. Indeed, in contrast to the effects of four days of L-NAME pretreatment, 10-minute pretreatment with L-NAME significantly reduce the duration of vasodilation mediated by bolus injection of alb-heme-NO, further supporting the notion that alb-heme-NO is a nitrodilator.

### Chemical evidence for the involvement of NANOS in heme-NO-mediated vasodilation

In addition to functional evidence, this study also provides chemical evidence that heme-NO mobilizes NO moieties from the NANOS within the arterial wall. Our experiments with ^15^N-tagged nitrite demonstrated that incubation with alb-heme-NO decreased arterial wall concentrations of a nitrogen oxide detected as ^15^N-nitrate while increasing ^15^N-NOx concentrations in the abluminal buffer, indicating an efflux of nitrite-derived NOx from the arteries into the buffer. Such an efflux has previously been observed with NTG, GSNO, and UV light^2,8^, suggesting a shared involvement of the NANOS in the vasodilation mediated by alb-heme-NO, GSNO, NTG, and UV light. Recent experiments have identified the arterial NANOS as a high molecular weight compound detected as nitrate^8^. The chemical nature of the NANOS, along with other components of the NANOS working model—such as the membrane receptor for nitrodilators and the mechanisms involved in the uptake, retention, and release of NO bioactivity^2^— will be subjects of our future studies. It is important to emphasize that while the current study suggests heme-NO does not cross the plasma membrane for signaling, it does not rule out heme-NO as an intracellular signaling entity. In fact, as a mobile and exchangeable NO escort, heme-NO offers a promising route for regulating the intracellular NO moieties in the NANOS.

### Role of heme-NO in the export of NO bioactivity from RBCs

Blood and RBCs have long been recognized for their ability to transport NO bioactivity in an endocrine manner^32,33^. Various NO species, including SNO, nitrite, and more recently heme-NO, have been proposed as intermediates in this process^3,32,33^. Notably, heme-NO emerges as a particularly compelling candidate, as it resists scavenging reactions^3,6,7^ and selectively activates sGC with the same efficacy as NO^4,5^. The potential vasoactive role of heme-NO is supported by the formation of HbNO in the blood of humans undergoing NO inhalation or nitrite administration, as well as by the observed artery-to-vein gradients of HbNO during NO inhalation in humans^27,34^. In line with this, our studies suggested that the placenta generates heme-NO, likely in the form of HbNO, from nitrite, and that a vein-to-artery gradient of heme-NO exists in the umbilical cord^35–37^. Nevertheless, the current study found that purified free HbNO does not exhibit vasodilatory effects in isolated arteries. Additionally, during the room temperature storage of deoxygenated blood pretreated with NO to form HbNO, its vasodilatory activity diminishes over the course of an hour despite unchanged levels of heme-NO and HbNO. In line with the absence of an efficient mechanism for the export of heme or heme-NO from RBCs, these findings challenge the role of heme-NO derived from HbNO as the intermediate for exporting NO bioactivity from RBCs.

In conclusion, we found that heme-NO does not cross the plasma membrane for signaling; instead, it dilates arteries by mobilizing NO moieties from a vascular intracellular NO store.

## Declaration of competing interest

Drs. Liu and Blood disclose they are named on a patent for the combined use of nitrite and nitrodilators for cardiovascular therapeutic use.

## Funding

The studies were supported by NIH grants HL095973 (ABB), HD083132 (LZ), and HL155295 (ABB/LZ).

## Acknowledgements

The authors appreciate input of Dr. Hobe Schroeder (Loma Linda University) via helpful discussions and Linda Liu (Cope Middle School, Redlands, CA) for artwork. Daniel Castella is acknowledging support from a University of Michigan Rackham Merit Fellowship.

